# AtlasXplore: a web platform for visualizing and sharing spatial epigenome data

**DOI:** 10.1101/2023.04.23.537969

**Authors:** Joshua Barnett, Noori Sotudeh, Poorvi Rao, Jonah Silverman, Tamara Jafar, Liya Wang

## Abstract

In recent years, a growing number of spatial epigenome datasets have been generated, presenting rich opportunities for studying the regulation mechanisms in solid tissue sections. However, visual exploration of these datasets requires extensive computational processing of raw data, presenting a challenge for researchers without advanced computational skills to fully explore and analyze such datasets. Here we introduce AtlasXplore™, a web-based platform that enables scientists to interactively navigate a growing collection of spatial epigenome data using an expanding set of tools.

**Availability and implementation:** AtlasXplore is freely available at https://web.atlasxomics.com

## 1 Background

The rapid development of single-cell transcriptomics (scRNAseq) and epigenomics (scATACseq, scCUT&Tag) in the last decade has unraveled the composition of cell types isolated from dissolved tissues as well as the interaction between transcriptional machinery and chromatin states. However, single-cell technologies have intrinsic limitations in studying the interactions between cells and their microenvironment, as spatial information is lost during tissue dissociation. The last few years have seen the development of multiple spatial transcriptomics methods, used to assay transcriptomes while keeping the tissue sections from which they came intact (Moses and Pachter, 2022). Recently, spatial epigenome assays have emerged for interrogating chromatin state (Deng, Bartosovic, Ma, *et al*., 2022; Llorens-Bobadilla *et al*., 2023), histone modifications (Deng, Bartosovic, Kukanja, *et al*., 2022), or co-profiling chromatin state/histone modification with transcriptome (Zhang *et al*., 2023) in solid tissue sections. These new assays create a need for interactive tools to facilitate the exploration and sharing of such high-dimensional spatial epigenome datasets.

Following alignment and counting of the raw data (generating the fragment file) and image data analysis (Barnett et al., 2022), researchers typically proceed with the computation of quality control metrics, clustering and embedding, marker identification, peak calling, motif searching, generation of genome browser tracks, scRNAseq data integration (if available), regulation network analysis (if co-profiled with transcriptome), and projections of various scores onto the tissue images. AtlasXplore integrates the multiple layers of epigenome data into a web browser-based interactive viewer.

## 2 Implementation

The architecture of AtlasXplore’s application is illustrated in Figure 1A. In addition to visualization, Celery (with RabbitMQ and redis) is used for queuing asynchronous tasks, such as cell type identification with user-provided markers, identifying the top ten features in a lasso selection, injecting spatial data into the platform, and subsetting the regulation network. Results are returned to the front end (Vue.js) and displayed.

**Figure 1.**
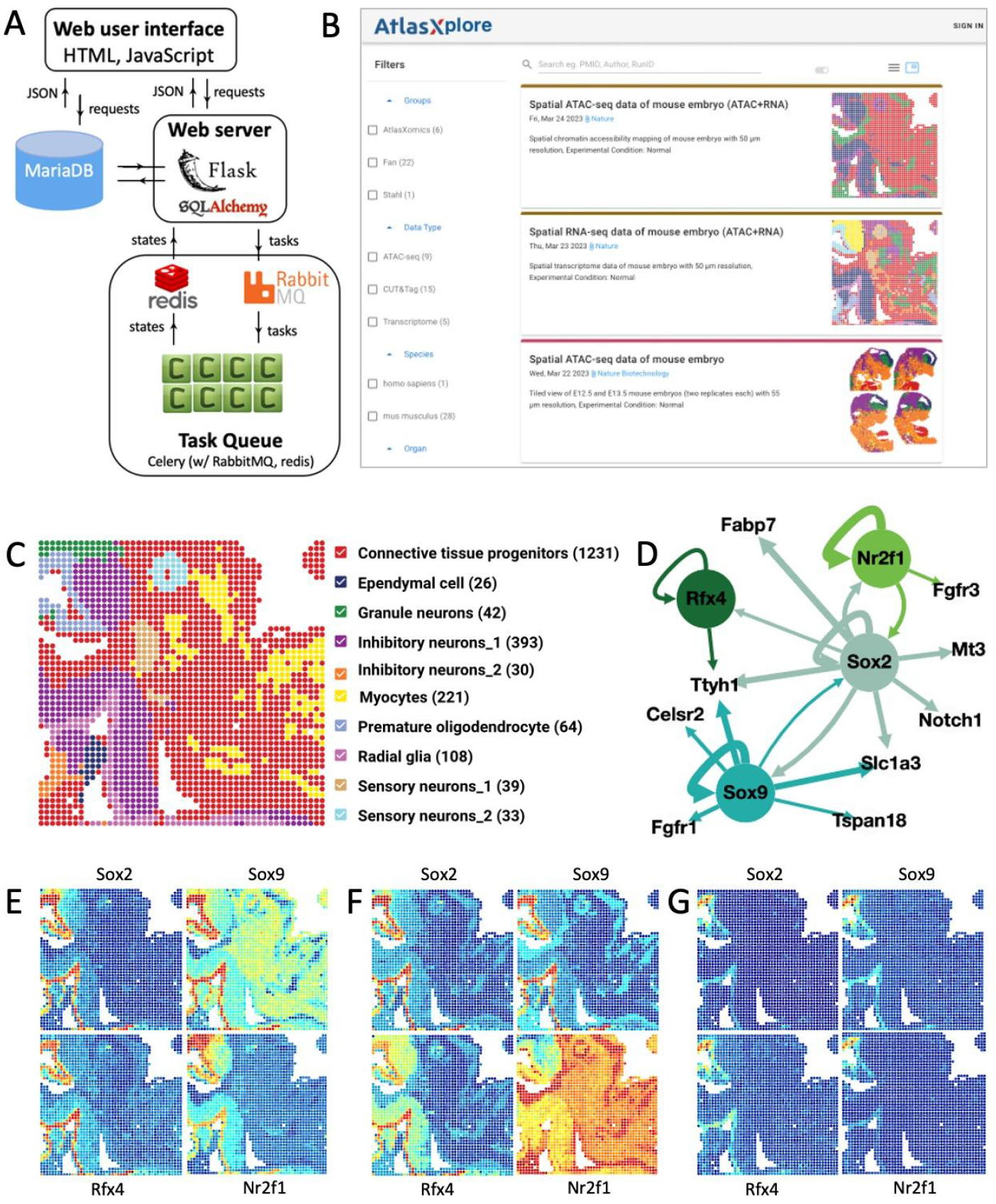
A. Software architecture; B. Landing page; C. Cell types projected on the tissue for E13 mouse embryo; D. Regulation network built from co-profiling data; E. Accessibility scores; F. Motif deviation scores; G. AUC score of 4 TFs.

AtlasXplore protects private data with Amazon Cognito authentication, and makes published and exemplar data accessible to both guests and registered users. Users can search via PMID/author or filter by research group, type, species, and tissue (Figure 1B).

## 3 Features

AtlasXplore supports three modalities of interactive exploration: gene, motif, and eRegulon (González-Blas *et al*., 2022). There are three panels used to display each data type. The top panels have the spatial map (left) and UMAP (right) to display the activity score (Granja *et al*., 2021) for genes assayed with ATAC or CUT&Tag active epigenetic antibodies such as H3K4me3, chromatin silencing score (Wu *et al*., 2021) for genes with CUT&Tag repressive epigenetic mark like H3K27me3, as well as deviation score for motifs and AUC score (Aibar *et al*., 2017) for eRegulons. The bottom panel displays the top up/down regulated features per cluster which can be toggled to display the histogram view per feature, the genome browser view per gene (Figure S1), the seqLogo plot per motif, or the regulation network for one or more eRegulons (Figure 1D). The search box on the top (with autocomplete) is the placeholder for checking a single or a collection of genes/motifs/eRegulons. The button next to the search bar toggles between displaying the mean or individual views. For example, the individual view of 4 transcription factors (TFs) is shown for the activity score (Figure 1E), motif deviation score (Figure 1F), and eRegulon AUC score (Figure 1G).

As mentioned in the last section, Celery workers are used to compute top 10 features from lasso selections performed on either UMAP or spatial map. Spot indices and top 10 features inside the selection are returned for analysis. For example, the top 10 features can be pasted back to the search box for visual inspection. These workers are also used to support cell type mapping where a statistical t test is implemented to assign each cluster to a cell type with a list of markers for each cell type provided by users. For simplifying the visualization of the regulation network, Celery workers are used to retrieve the subnetwork with given TFs. The subnetwork can be simplified by keeping only top regulations at the level of 15%, 25%, 50%, or 75%.

## 4 Advanced features demonstrated with the spatial co-profiling data

Following the Scenic+ procedure (González-Blas *et al*., 2022), the co-profiled E13 mouse embryo data (Zhang *et al*., 2023) are processed to build the regulation network. Cell types are identified with the reference MOCA RNAseq data (Cao *et al*., 2019) and refined with unsupervised clusters and spatial data. Figure 1C shows the cell types projected to the tissue. Each cell type is color-coded and the number of spots per cell type is counted. The resulting network contains more than 100 eRegulons which can be subsetted with TFs via search. As an example, the subnetwork of Sox2, a well known Radial glia (RG) marker (Li *et al*., 2011), is shown in Figure 1D. The subnetwork is retrieved by keeping only genes and TFs regulated by Sox2, and it is simplified with the top 25% significant regulations. In the network, TFs are color-coded, and the thickness of the connection is proportional to the regulation strength.

The promoters of all 4 TFs in the network are highly accessible in RG and Premature oligodendrocyte (PO) cells (Figure 1E), which is not surprising since their expressions are regulated by themselves and also other TFs. The binding site accessibilities of these TFs are highly increased in RG and PO cells, as demonstrated by their motif deviation score (Figure 1F). This suggests that these TFs could be critical regulation players in these two cell types. Therefore the AUC scores of all 4 eRegulons are significantly higher in RG and PO cells (Figure 1G). These are consistent with earlier findings: Sox9 regulates the generation of astrocytes and oligodendrocytes from glial precursor cells (Stolt and Wegner, 2010); Nr2f1 plays a key role in the maturation of oligodendrocytes (Tocco *et al*., 2021); and Rfx4 might play a role in the specification and development of glia (Zhang *et al*., 2006).

The subnetwork shows that Sox2 is regulated by 3 TFs (Nr2f1, Sox9, and itself), and there are three highly accessible regions/peaks before its promoter region, as shown in Figure S1. In addition, the accessibilities is significantly higher in cluster 5 (RG and PO cells) than other clusters.

## 5 Future work

As demonstrated, AtlasXplore integrates multiple layers of spatial epigenome data for deep diving into the biological insights buried inside the data. With the integration with Celery workers, there is unlimited potential for AtlasXplore to incorporate other software and functions for interactive exploration of high-dimensional data sets. One ongoing project is to extend multiple sample comparisons (e.g. per cluster pairwise comparisons) after tiling them together (as the 4 replicates of mouse embryos shown in Figure 1B). Another ongoing project is packaging AtlasXplore as standalone software for running locally. Currently, AtlasXplore serves as a repository for emerging spatial epigenome data.

---

**Figure S1.**
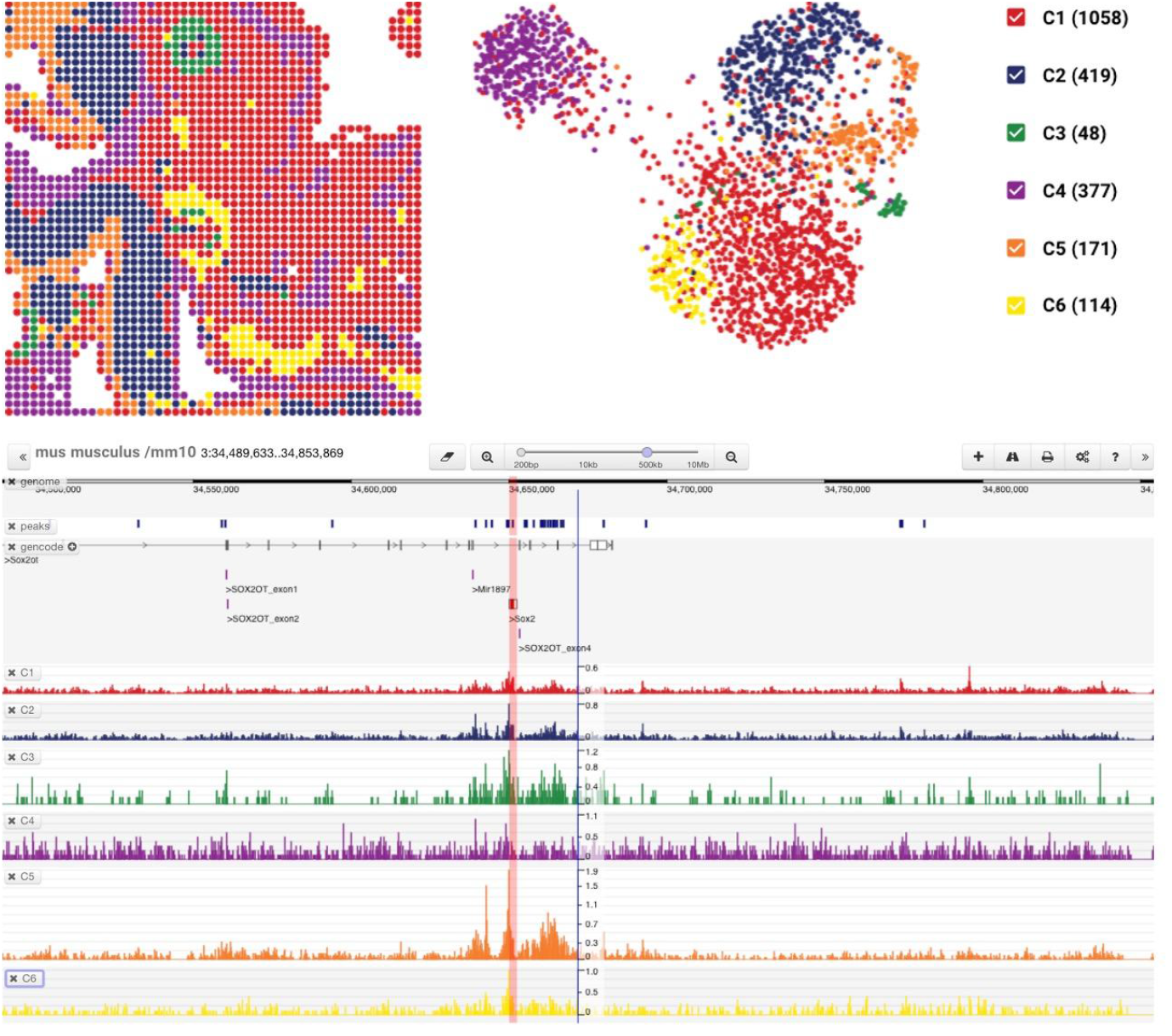
Spatial and Umap plots (top) and genome browser view of Sox2 (bottom) for the spatial ATAC mouse embryo data.

